# TemStaPro: protein thermostability prediction using sequence representations from protein language models

**DOI:** 10.1101/2023.03.27.534365

**Authors:** Ieva Pudžiuvelytė, Kliment Olechnovič, Egle Godliauskaite, Kristupas Sermokas, Tomas Urbaitis, Giedrius Gasiunas, Darius Kazlauskas

## Abstract

**Motivation:** Reliable prediction of protein thermostability from its sequence is valuable for both academic and industrial research. This prediction problem can be tackled using machine learning and by taking advantage of the recent blossoming of deep learning methods for sequence analysis. These methods can facilitate training on more data and, possibly, enable development of more versatile thermostability predictors for multiple ranges of temperatures.

**Results:** We applied the principle of transfer learning to predict protein thermostability using embeddings generated by protein language models (pLMs) from an input protein sequence. We used large pLMs that were pre-trained on hundreds of millions of known sequences. The embeddings from such models allowed us to efficiently train and validate a high-performing prediction method using over one million sequences that we collected from organisms with annotated growth temperatures. Our method, TemStaPro (Temperatures of Stability for Proteins), was used to predict thermostability of CRISPR-Cas Class II effector proteins (C2EPs). Predictions indicated sharp differences among groups of C2EPs in terms of thermostability and were largely in tune with previously published and our newly obtained experimental data.

**Availability and Implementation:** TemStaPro software and the related data are freely available from https://github.com/ievapudz/TemStaPro and https://doi.org/10.5281/zenodo.7743637.

## 1. Introduction

Biotechnological research and development often involves searching for proteins that can remain stable (maintain their spatial structures) in a high-temperature setting. In many cases, the only information initially known about a protein is its sequence of amino acids. Therefore it is beneficial to have computational tools that can efficiently predict protein thermostability from a protein sequence alone. Several machine learning-based methods were developed for that in the past (Gromiha and Suresh, 2008; Lin and Chen, 2011; Fan et al., 2016) and in recent years (Feng et al., 2020; Charoenkwan et al., 2021, 2022; Ahmed et al., 2022; Zhao et al., 2023), but these efforts did not focus on drastically increasing the amount of data used for training and validation, which could potentially lead to better-performing methods with an ability to distinguish multiple levels of thermostability.

In general, most of the current state-of-the-art sequence-based methods (Feng et al., 2020; Charoenkwan et al., 2021, 2022; Ahmed et al., 2022) were trained and tested using protein sequences taken from datasets of proteins annotated with experimentally-determined thermal stability information. Such datasets are inevitably small because gathering experimental data on a per-protein basis is usually very expensive and time-consuming. There is an alternative way of collecting protein thermostability data — taking the available information about optimal growth temperatures of organisms that have sequenced genomes converted to proteomes, grouping the proteomes by the corresponding growth temperature intervals, collecting the protein sequences, and annotating them with temperature values (Engqvist, 2018a; Zhao et al., 2023). This approach is not as precise as gathering experimental data for every protein separately, but the optimal growth temperature of an organism provides a reliable lower bound for the melting temperature of proteins in that organism (Dehouck et al., 2008). Most importantly, the proteomes-based data gathering can provide millions of sequences for machine learning. Nevertheless, even one of the most recent deep learning-based methods (Zhao et al., 2023) that used such data was trained only on a small subset (less than 1%) of sequences from available proteomes with known growth temperatures.

Training using big data when starting from raw amino acid sequences is extremely challenging due to the need to construct or learn a complex protein representation suitable for making predictions. However, there is a possibility to take a shortcut and apply a transfer learning approach — use protein representations generated by other methods trained for different tasks. More specifically, it is possible to use protein sequence embeddings generated by encoders of protein language models (pLMs) that were trained on hundreds of millions of natural protein sequences, e.g. ESM (Rives et al., 2021; Lin et al., 2023) and ProtTrans (Elnaggar et al., 2022). Such pLM embeddings are rich representations that were already shown to be suitable inputs for various predictive tasks (Fenoy et al., 2022). It was only a matter of time before pLM embeddings were also applied for the identification of thermostable proteins. To the best of our knowledge, BertThermo (Pei et al., 2023) was the first such method that was published. However, BertThermo was trained using only 2803 sequences and was based on a binary classifier for only a single temperature threshold. Another recently published method, ProLaTherm (Haselbeck et al., 2023), was not substantially superior in this regard, it used only 7409 sequences split into just two classes of thermostability.

In this work we propose a large-scale comprehensive approach to using pLM embeddings for predicting whether a protein remains stable above some temperature threshold. We collected over one million of protein sequences from organisms with known optimal growth temperatures and we used that data to train, validate, and test multiple binary classifiers for multiple temperature thresholds. We showed that our classifiers perform exceedingly well both on our newly introduced test sets and on previously published benchmark datasets. We combined the classifiers into a software tool that, given a protein sequence as input, predicts protein stability for multiple temperature thresholds and checks if the predictions are not contradicting each other. The resulting method, *TemStaPro* (Temperatures of Stability for Proteins), is freely available as a standalone program.

We tested TemStaPro software to predict thermostability of CRISPR-Cas Class II effector proteins (C2EPs). C2EPs are usually found in bacteria that grow best in moderate temperatures (20 to 45 °C). However, there are few Cas9 and Cas12b variants that can function at temperatures above 60 °C (Gasiunas et al., 2020; Nguyen et al., 2021). Thermostable C2EPs are important because they can be: used in conjunction with nucleic acid amplification methods to detect SARS-CoV-2 variants of concern in a single reaction (Nguyen et al., 2021); used in other CRISPR-based diagnostics methods (Ghouneimy et al., 2023); used for genome engineering of thermophilic organisms (Adalsteinsson et al., 2021); used in aid to increase lifetime of gene editing tools in human plasma (Harrington et al., 2017). Our results indicate that thermostability differs among groups of C2EPs, for example Cas12f and TnpB-like proteins are more likely to function at higher temperatures than ones from Cas9 and Cas13 groups.

## 2. Materials and methods

### 2.1 Data preparation

The data source that was used to construct training, validation, and testing datasets was composed of 21 498 annotated organisms (Engqvist, 2018b,a). The taxonomy identifiers given in the data source were used to fetch UniParc (Leinonen et al., 2004) identifiers for the corresponding proteome. UniParc identifiers were used to download FASTA files with proteins composing the proteomes. The set of proteomes was filtered so that only non-eukaryotic proteomes with non-duplicated taxonomy identifiers remained.

The sequences of the final 5491 proteomes were clustered with CD-HIT (Fu et al., 2012) using 30% sequence identity threshold. The clustering results were later utilized to ensure that the training, validation, and testing sets (employed for machine learning) would always consist of sequences from different clusters, preventing any overlap of sequences within the same cluster across such sets.

The collection of datasets that were used to train, validate, and test the tool is summarized in Table 1. The collected and clustered proteomic sequences were used to construct two different-sized datasets for machine learning, *TemStaPro-Minor-30* (derived from randomly selected 162 proteomes) and *TemStaPro-Major-30* (derived from all downloaded proteomes).

**Table 1.** Datasets of protein sequences that were used in the development of the tool.

| Dataset | Subset | All sequences | Class 0 sequences | Class 1 sequences | Max. length |
| --- | --- | --- | --- | --- | --- |
| TemStaPro-Minor-30 | Cross-validation | 239 629 | 146 657 | 92 972 | 1 022 |
|  | Testing | 41 930 | 25 580 | 16 350 | 1 022 |
| TemStaPro-Major-30 | Training | 943 605 | 879 799 | 63 806 | 9 841 |
|  | Validation | 230 985 | 215 167 | 15 818 | 9 387 |
|  | Testing | 210 137 | 196 072 | 14 065 | 13 394 |
| SAPPHERE | Testing | 742 | 371 | 371 | 1 643 |
| iThermo | Testing | 562 | 289 | 273 | 3 567 |

The smaller dataset, *TemStaPro-Minor-30*, was designated for the single-threshold binary classifier design process in a 10-fold cross-validation setting. The smaller data size was crucial to rapidly iterate through different inputs as well as different machine learning model architectures and hyperparameters. To be processable with length-limited pLMs such as ESM-1b (Rives et al., 2021), the dataset was restricted to sequences no longer than 1022 amino acids. With *TemStaPro-Minor-30*, the main objective was to design a classifier for proteins that are stable (class ‘1’) or not stable (class ‘0’) at 65 °C and higher temperatures. The threshold of 65 °C was chosen because primary envisioned purpose of our method was to detect C2EP proteins that can withstand *>*60 °C temperatures during isothermal amplification step in a one-pot SARS-CoV-2 detection kit (Nguyen et al., 2022). Also, *TemStaPro-Minor-30* is hardly suitable for training for multiple thresholds because, due to the random sampling of proteomes, it contains protein sequences mostly from organisms, whose growth temperatures are in ranges 20-40 °C or 60-80 °C (Supplementary Fig. S1), thus the cases, which are not included in these intervals, are not sufficiently covered.

The larger dataset, *TemStaPro-Major-30*, was designated to involve as much data as possible to train, validate and test binary classifiers for 9 different temperature thresholds (40, 45, 50, 55, 60, 65, 70, 75, 80 °C) for the final software tool, utilizing the input and the architecture selected during the classifier design process performed using *TemStaPro-Minor-30*. Importantly, *TemStaPro-Major-30* sequences cover the range of temperatures between 0 and 100 °C (Supplementary Fig. S2).

It is important to note that the collected data is not suitable for training multi-class classification (where classes represent non-overlapping temperature intervals) or regression models because the temperature values used for ground truth are only lower bounds for the possible temperatures of stability of proteins. For example, a protein from an organism living in 45 °C environment may (or may not) also be stable at 60 °C. This also means that a single binary classifier is not very versatile. A binary classifier trained using 45 °C threshold cannot tell if a protein predicted as stable at over 45 °C is also stable at over 60 °C — a classifier trained using at least 60 °C threshold is needed for that.

### 2.2 Additional data for benchmarking

#### 2.2.1. SAPPHIRE and iThermo testing sets

SAPPHIRE (Charoenkwan et al., 2022) dataset was used to compare the performance of TemStaPro with SAPPHIRE, SCMTPP, and ThermoPred methods. This dataset is a balanced set of 742 sequences. It contains thermophilic and non-thermophilic proteins — the exact temperature threshold for group distinction is not mentioned, but, according to the creators of the dataset, proteins labelled as thermophilic are stable at 80-100 °C temperatures, they were taken from thermophilic organisms that grow at those temperatures (Charoenkwan et al., 2021).

iThermo (Ahmed et al., 2022) dataset was used to compare the performance of TemStaPro with iThermo, BertThermo, and DeepTP methods. This dataset is composed of proteins from organisms that grow in vastly different temperature ranges: above 60 °C and below 30 °C.

#### 2.2.2. Sequence dataset of Class II effector proteins

Initial datasets of Cas12 and Cas9 were taken from (Sasnauskas et al., 2023) and (Gasiunas et al., 2020), respectively. Cas13 sequence dataset was constructed by building HMMER (Eddy, 2011) sequence profiles for Cas13 groups (Kavuri et al., 2022) and using them to search NR (Sayers et al., 2022), UniRef100 (Suzek et al., 2007), MGnify (Richardson et al., 2023), and IMG/VR v4 (Camargo et al., 2023) databases with hmmsearch (Eddy, 2011). Only sequences with E-value ≤1e-20 were extracted. In case the same sequence was found using different queries, the hit having lower E-value was assigned to the group. Latter dataset was combined with Cas12 and Cas9 datasets to form final dataset of 16376 sequences (*SupplementaryFileC2EPsPredictions*.*tsv*). Thermostability predictions were done for all those sequences, but to check the thermostability of different C2EP groups, we only used sequences devoid of prediction clashes and having more than 300 residues.

### 2.3 Protein language models

This work exploits the transfer learning by taking protein representations from the last layer of protein language models and passing them as input to the classification model (Fig. 1).

**Fig. 1:**
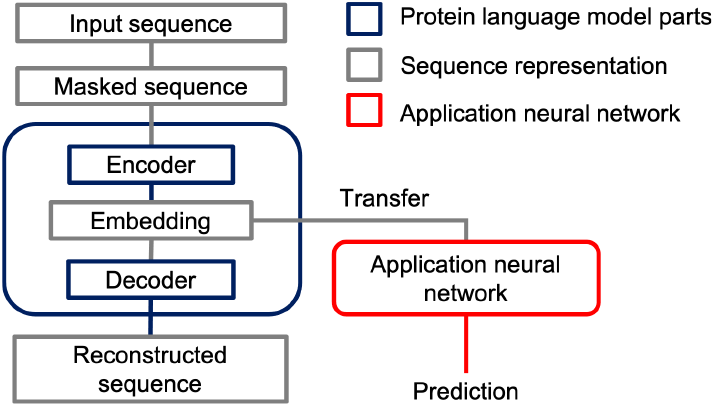
The scheme of embeddings from protein language model usage in the application neural network model.

The potential suitability of both ESM-2 and ProtT5-XL embeddings for thermostability classification was detected using principal component analysis (PCA) of mean embedding vectors. Plots of the first two principal components (Fig. 2) demonstrated the distinct separation of points corresponding to different thermostability classes.

**Fig. 2:**
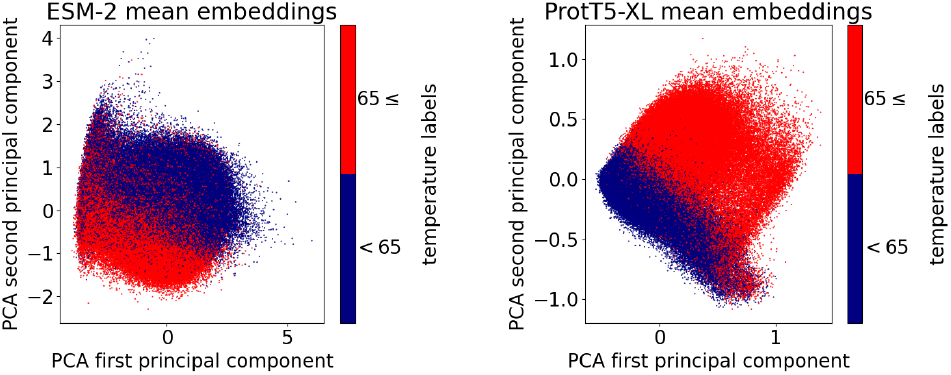
PCA visualizations of ESM-2 mean embeddings (left) and ProtTrans ProtT5-XL mean embeddings (right) computed for the cross-validation subset of the *TemStaPro-Minor-30* dataset. Plots were generated using *Scikit-Learn Python* library (version 1.3.2).

### 2.4 Binary classifier design process

The classifier design process utilized *TemStaPro-Minor-30* dataset and involved 10-fold cross-validation and testing of multiple neural network (NN) architectures using different pLM embeddings (ESM-2 or ProtTrans ProtT5-XL). Classifiers in this study were implemented as feed-forward densely connected NN models with up to two hidden layers, whose sizes were chosen to be original embeddings size divided by several multiples of 2 (Supplementary Fig. S3, Supplementary Table S1). The performance analysis was done using common metrics for binary classification (Matthews correlation coefficient (MCC), accuracy, precision, recall, and area under the receiver operating characteristic curve (ROC AUC)), with MCC being the primary metric.

The cross-validation results showed that all the tried combinations of inputs and NN architectures with at least one hidden layer achieve very similar high MCC values (Supplementary Fig. S4) Usage of a non-zero weight decay had a slight effect on the performance results (Supplementary Fig. S5, S6).

The results of testing performed using model trained on the whole unsplit cross-validation (using the average stopping epoch learned from the cross-validation stage) were highly consistent with the cross-validation results (Supplementary Fig. S7).

Overall, ESM-2-based and ProtT5-XL-based variants performed very similarly, yet ProtT5-XL embeddings are faster to compute and shorter, therefore the best architecture utilizing ProtT5-XL was chosen (C2H2 h256-128 from Supplementary Table S1).

### 2.5 Final binary classifiers training and testing

The functionality of the TemStaPro method is carried out by multiple binary classifiers that accept mean ProtT5-XL (from ProtTrans) representations (vectors of length 1024) as input. Classifiers were implemented as NN models using *PyTorch*. Each model is a multi-layer perceptron with 2 fully-connected hidden layers of sizes 256 and 128. After each layer (except the last one), rectified linear unit (ReLU) activation function is applied. Sigmoid activation function is used after the last layer.

Every single binary classifier was trained on *TemStaPro-Major-30* data (labelled according to the corresponding temperature threshold), using mini-batch training principle (with batch size of 24) and Adam optimizer (Kingma and Ba, 2017) with learning rate of 0.0001 and weight decay of 0.0001. Since the datasets were imbalanced, training and validation sets were loaded using weighted random sampler.

Each binary classifier was validated by assessing performance on the whole validation set after every training epoch, selecting the model that achieved the highest MCC. Each chosen model was then tested on the appropriate testing set, reporting MCC and other binary classification metrics.

### 2.6 Predictor application

The TemStaPro user’s input is a FASTA file with amino acid sequences with proteins’ identifiers in the headers. For each protein in the FASTA file a mean ProtT5-XL (Elnaggar et al., 2022) embedding is generated, which is the input of the classification model. In addition to this, there is an option available to pass embeddings of each residue to the model. Per-residue embeddings can also be averaged over a residue window of size k, which can be customized, to get per-segment embeddings. Then each segment (of size k) of amino acids gets a prediction.

In the default TemStaPro operating mode, the classification predictions are made by 6 ensembles each composed of 5 neural network models that were trained to make binary classification of proteins with respect to one of 6 temperature thresholds: 40, 45, 50, 55, 60, and 65 °C. The output list of 6 predictions is created from averaged predictions of each ensemble.

Based on the sequence of all 6 classification predictions, each input protein is assigned two labels: left-hand and right-hand. These labels are determined by scanning the binary predictions starting from the left or right-hand side, respectively. For example, the left-hand label is assigned the temperature range, where the last positive prediction (class ‘1’) is encountered. If outputs are only negative (class ‘0’), then the label is the first temperature range. On the other hand, the right-hand label is assigned by reading the outputs starting from the right: the label is assigned the temperature range, where the first ‘1’ is encountered. The treatment of the ‘0’-only case coincides with the left-hand principle.

Since binary predictions are made independently, conflicts might occur between the outputs of the classifiers: for instance, the predictor of 40 °C threshold would predict that protein is not stable at 40 °C and higher temperatures, although the predictor of 50 °C threshold would state otherwise. When such conflict occurs, left-hand and right-hand labels differ. On the contrary, if the labels report the same temperature interval, that interval can be interpreted as the highest temperature range at which the protein was predicted to still be thermostable.

An alternative TemStaPro operating mode is similar to the default one, but includes three more classifiers (for thresholds of 70, 75, and 80 °C). Due to relative scarcity of positive examples for training these additional classifiers, they are expected to be less reliable and therefore they were not included in the default TemStaPro mode.

## 3. Results

### 3.1 Performance of binary classifiers

Performance of the trained classifiers on the *TemStaPro-Major-30* testing set is summarized in Supplementary Figure S8 and Table S2. The maximum MCC score was 0.638, achieved by the binary classifier for 50 °C temperature threshold. The MCC scores for the classifiers of the highest practical utility were: 0.601 for 60 °C, 0.610 for 65 °C.

However, the performance metrics may be not easy to interpret as they are affected by the substantial threshold-dependent imbalance of positive and negative examples in the *TemStaPro-Major-30* testing set. A balanced testing set may provide a more conventionally interpretable performance overview. It is also important to assess results in comparison with other methods that use pLM embeddings. Thus, for each threshold, a balanced testing subset of 2000 sequences was randomly sampled from *TemStaPro-Major-30*. The sampled data was used to compare performance of BertThermo, ProLaTherm, and TemStaPro for every threshold. The resulting MCC values are showcased in Fig. 3, other metrics are presented in Supplementary Table S3. For all the temperature thresholds, TemStaPro classifiers reached higher MCC, PR AUC, ROC AUC, F1, and accuracy scores than BertThermo and ProLaTherm. The maximum MCC score was 0.903, achieved by the TemStaPro binary classifier for 80 °C temperature threshold. The MCC scores for the TemStaPro classifiers of highest practical utility were: 0.805 for 60 °C, 0.811 for 65 °C.

**Fig. 3:**
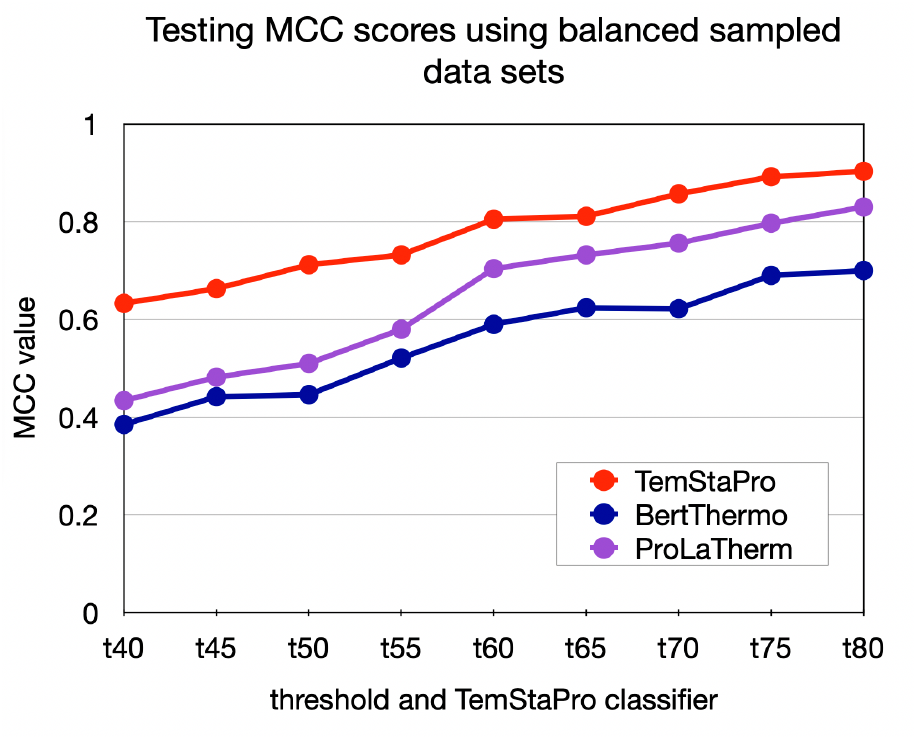
Comparison of testing scores between BertThermo, ProLaTherm, and TemStaPro using the sampled balanced testing datasets for each threshold included in TemStaPro.

### 3.2 Performance on other benchmarks

To compare TemStaPro with modern, yet non-deep learning-based methods, TemStaPro was tested on the SAPPHIRE tool’s testing dataset (Charoenkwan et al., 2022), which was previously used to test SCMTPP (Charoenkwan et al., 2021) and ThermoPred (Lin and Chen, 2011) tools for protein thermostability prediction.

The method with the highest MCC score among SCMTPP, ThermoPred, and SAPPHIRE is the latter one. The results showed that TemStaPro predictors for temperature thresholds 60-70 °C perform better than SAPPHIRE (Table 2). The MCC scores of our other predictors (trained for thresholds ≤ 55 and ≥ 75 ^°^C) differed from SAPPHIRE tool’s score by no more than 0.08.

**Table 2.** Models’ scores after testing with an independent SAPPHIRE dataset. Each column’s highest values are emphasized in bold.

| Model | Accuracy | Sensi-<br>tivity | Spe-<br>cificity | MCC | ROC<br>AUC |
| --- | --- | --- | --- | --- | --- |
| TemStaPro-t80 | 0.923 | 0.849 | **0.997** | 0.856 | 0.990 |
| TemStaPro-t75 | 0.930 | 0.868 | 0.992 | 0.867 | 0.989 |
| TemStaPro-t70 | **0.960** | 0.951 | 0.968 | **0.919** | 0.993 |
| TemStaPro-t65 | 0.957 | 0.957 | 0.957 | 0.914 | 0.993 |
| TemStaPro-t60 | 0.951 | 0.987 | 0.916 | 0.905 | 0.995 |
| TemStaPro-t55 | 0.934 | 0.989 | 0.879 | 0.873 | **0.996** |
| TemStaPro-t50 | 0.908 | 0.989 | 0.827 | 0.828 | 0.993 |
| TemStaPro-t45 | 0.907 | 0.989 | 0.825 | 0.825 | 0.992 |
| TemStaPro-t40 | 0.896 | **0.995** | 0.798 | 0.808 | 0.993 |
| SAPHIRE | 0.942 | 0.951 | 0.933 | 0.884 | 0.980 |
| SCMTPP | 0.865 | 0.849 | 0.881 | 0.731 | - |
| ThermoPred | 0.86 | 0.938 | 0.782 | 0.729 | - |

To compare TemStaPro with modern deep learning-based methods, it was tested on the iThermo (Ahmed et al., 2022) test dataset. The DeepTP (Zhao et al., 2023) and BertThermo (Pei et al., 2023) performance results were taken from the BertThermo publication. Among BertThermo, DeepTP, and iThermo, BertThermo achieved the best evaluation classification scores. TemStaPro outperformed BertThermo — the maximums of all the metrics were achieved by the TemStaPro classifiers (Table 3).

**Table 3.** Models’ scores after testing with iThermo dataset. Each column’s highest values are emphasized in bold.

| Model | Accuracy | Sensi-<br>tivity | Spe-<br>cificity | MCC | ROC<br>AUC |
| --- | --- | --- | --- | --- | --- |
| TemStaPro-t80 | 0.884 | 0.766 | **0.997** | 0.787 | 0.986 |
| TemStaPro-t75 | 0.902 | 0.802 | **0.997** | 0.818 | 0.990 |
| TemStaPro-t70 | 0.973 | 0.952 | 0.993 | 0.947 | **0.999** |
| TemStaPro-t65 | **0.986** | 0.978 | 0.993 | **0.972** | 0.998 |
| TemStaPro-t60 | 0.979 | 0.989 | 0.969 | 0.958 | **0.999** |
| TemStaPro-t55 | 0.980 | **1.000** | 0.962 | 0.962 | **0.999** |
| TemStaPro-t50 | 0.963 | **1.000** | 0.927 | 0.928 | **0.999** |
| TemStaPro-t45 | 0.964 | **1.000** | 0.931 | 0.931 | **0.999** |
| TemStaPro-t40 | 0.954 | 0.996 | 0.913 | 0.911 | **0.999** |
| iThermo | 0.963 | 0.963 | 0.962 | 0.927 | 0.986 |
| DeepTP | 0.934 | 0.963 | 0.901 | 0.870 | 0.983 |
| BertThermo | 0.975 | 0.985 | 0.965 | 0.950 | 0.994 |

Importantly, TemStaPro training data was made to not contain any sequences closer than 30% in terms of identity to the sequences from SAPPHIRE and iThermo datasets.

### 3.3 Software tool

We implemented TemStaPro as a command-line software tool. By default, for each input protein sequence, TemStaPro outputs global predictions for 6 temperature thresholds, and the left-hand and right-hand labels derived from the predictions as described in section 2.6. An example output table of the global scoring is given in Supplementary Fig. S9.

Besides the default global protein scoring, the user might opt for per-residue or per-segment predictions. For the per-residue case each amino acid in the protein sequence gets a distinct set of thermostability predictions (Supplementary Fig. S10), similarly for per-segment option, where each full segment of the chosen size in the sequence gets its set of predictions. Additionally, there is an option to plot per-residue and per-segment predictions (Supplementary Fig. S11).

The speed of the TemStaPro software is mostly determined by whether the ProtT5-XL embeddings are produced on GPU or not. Using NVIDIA GeForce RTX 2080 Ti GPU, TemStaPro processes 10000 sequences with average length of 1000 residues in less than 2 hours. Without GPU, the operating time increases several times (up to 60 times if run on a laptop with Intel i7-8565U CPU).

### 3.4 Thermostability of Class II effector proteins

To get a better view on thermal stability among different C2EP groups, we tested our method on a large dataset (16376 sequences) of Cas9, Cas12, TnpB, and Cas13 proteins (*SupplementaryFileC2EPsPredictions*.*tsv*). For further analysis, we considered only sequences longer than 300 residues. There were 11341 such sequences, 10734 (94.6%) of them had clash-free predictions. Thermostability prediction varied greatly between groups of Cas12 and TnpB (Fig. 4). This might be explained by the fact that members of Cas12 and TnpB differ greatly in sequence similarity and length (Urbaitis et al., 2022; Karvelis et al., 2021). The Cas12a group, which is currently actively studied and used in biotechnology applications (Khan and Sallard, 2023), did not have thermostable (≥60 ^°^C) sequences (Fig. 4) as predicted by our method.

**Fig. 4:**
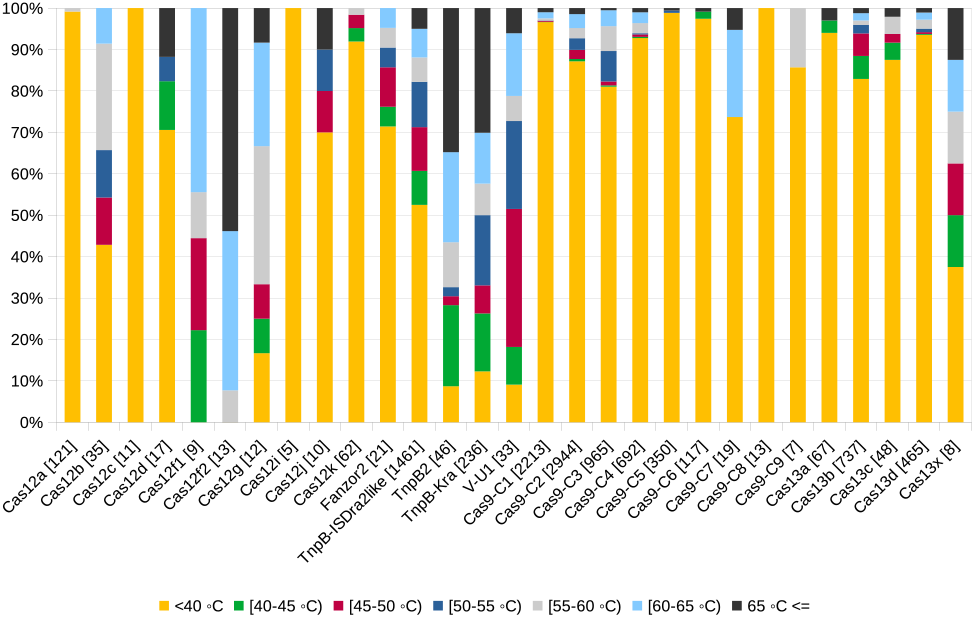
Predicted thermostability of various C2EP groups. Numbers in brackets correspond to the number of sequences tested.

In contrast to Cas12a, more than half of the members of Cas12b group were predicted to function at 45 °C or higher temperature (Fig. 4). Such observation corresponds to the experimental data because most of the characterized thermostable Cas12 proteins belong to the Cas12b group (Yang et al., 2016; Nguyen et al., 2022). Interestingly, we predicted that most thermostable Cas12 groups are Cas12f1, Cas12f2, and Cas12g (Fig. 4). However, the latter groups were not studied experimentally for thermostability. On the other hand, some of the members (e.g. Un1, Un2, Mi1, and Mi2) of Cas12f1 and Cas12f2 groups are found in archaea, which is an indication of possible thermostability of these C2EPs. Two groups of TnpBs (namely, TnpB2 and TnpB-Kra, which contains TnpB from *Ktedonobacter racemifer* (Altae-Tran et al., 2021)) showed higher predicted thermal stability compared to a group represented by TnpB from *Deinococcus radiodurans* ISDra2.

Just a few Cas9 groups (namely, Cas9-C3 and Cas9-C7; Supplementary Table S4) contain thermostable members. This observation is in tune with experimental data. The Cas9-C3 group contains characterized thermostable proteins CaldoCas9, GeoCas9, and ThermoCas9 (Adalsteinsson et al., 2021; Mougiakos et al., 2017; Harrington et al., 2017). Cas9-C7 group includes NsaCas9 which was shown to function at temperatures above 60 °C in our previous study (Gasiunas et al., 2020).

Cas13 groups tend to have predicted lower thermostability except for Cas13x (Fig. 4), which contains only 8 members, thus it is too early to draw any conclusions about their thermal stability.

In rare cases our method predicted lower (differences *>*10 °C) than experimentally characterized temperatures for thermostable proteins (e.g. TccCas13a; Supplementary Table S4). However, these are exceptions, in 90% of the cases (35 out of 39) predicted thermostability varied no more than 10 °C from the experimental data (Supplementary Table S4).

### 3.5 Experimental validation of thermostability predictions

To validate the accuracy of the thermostability prediction model, we have experimentally characterized two potentially thermostable proteins. Ghy2Cas9 was previously identified and described in (Gasiunas et al., 2020). This enzyme showed dsDNA cleavage activity in cell free lysates, but protein thermostability was not characterized. We also identified a putative thermostable Cas12b ortholog from *Clostridia bacterium*, CbaCas12b (NMA13999.1), in publicly available genome databases. We expressed the enzymes in *E. coli* and purified them. The proteins along with SpyCas9, as a control, were subjected to analysis by nano differential scanning fluorimetry (nanoDSF) to ascertain the temperatures at which they unfold. The enzymes were tested either without their guide RNA (apo) or with single guide RNA (RNP), except for CbaCas12b, for which we could not identify a tracrRNA. Ghy2Cas9 was shown to begin to unfold at around 54 °C, CbaCas12b at 57 °C, and SpyCas9 at 45 °C, with their respective RNPs unfolding at around 2-3 °C higher temperature (Supplementary figures S12a and S13). Predicted temperatures of thermal stability for both Ghy2Cas9 and SpyCas9 ([55-60) and *<*40 °C, respectively) did not differ more than 7 °C from their experimentally determined melting point temperatures. Following this, we evaluated the dsDNA cleavage activity of Ghy2Cas9 as well as SpyCas9 across a range of temperatures from 37 °C to 70 °C using fluorophore-labelled dsDNA substrates. As shown in Supplementary Fig. S12b, Ghy2Cas9 and SpyCas9 retained robust nuclease activity at temperatures up to 55 °C and 50 °C, respectively, which correlates with the determined unfolding temperature of the RNPs by nanoDSF.

## 4. Discussion and conclusions

Embeddings from pre-trained protein language models can be highly suitable for the task of protein thermostability prediction — this became nearly apparent even after our initial principal component analysis of ESM and ProtTrans embeddings. We further showed that a simple dense neural network can be efficiently trained to predict a protein thermal stability class from the mean of per-residue embedding vectors. With that established, we endeavored to make a better thermostability prediction method not by complicating the machine learning model, but rather by preparing and using more data for training and validation. We prepared and utilized a dataset of over one million sequences annotated with temperatures. The considerable amount and diversity of our data allowed us to train, validate, and test classifiers for multiple temperature thresholds (from 40 to 80 °C). For every temperature threshold, TemStaPro outperformed other pLM-based methods, BertThermo and ProLaTherm. When tested on recent independent datasets, SAPPHIRE and iThermo, our trained and validated method, named TemStaPro, performed better than state-of-the-art sequence-based predictors.

We combined our trained classifiers into a software tool that predicts protein thermostability for multiple temperature thresholds, reports whether the results of multiple classifiers are in agreement, and indicates the highest temperature range at which the protein is predicted to be thermostable. We tested that software on CRISPR-Cas Class II effector proteins. Interestingly, we saw large variation in thermal stability among groups of Cas12 and TnpB. For example, more than a half of the members of groups Cas12b, Cas12f1, Cas12f2, Cas12g, TnpB2, TnpB-Kra and V-U1 have predicted temperatures of ≥45 ^°^C (Fig. 4). In contrast, members of Cas12a, Cas9, and Cas13 groups might function at lower temperatures.

We also observed that TemStaPro is a more pessimistic than optimistic predictor — it tends to slightly underestimate the highest temperature at which the protein is still stable. We attribute this trait to the particularity of the training data, where sequences were annotated not with exact melting temperature values, but with their lower bounds.

To conclude, considering that the large majority (90%) of our predictions for well-characterized proteins were confirmed experimentally, we believe that TemStaPro can be useful for pre-screening potentially thermostable candidate proteins and thus reducing the number of experiments needed to determine protein thermostability in biotechnology.

## Data availability

The data underlying this article and a supplementary *\**.*tsv* table are available in Zenodo system at https://doi.org/10.5281/zenodo.7743637 Supplementary data *\**.*xlsx* tables are available at https://github.com/ievapudz/TemStaPro/tree/main/data.

TemStaPro software source code is freely available at https://github.com/ievapudz/TemStaPro.

## Acknowledgements

We thank Antanas Kiziela, Mindaugas Margelevičius, and Česlovas Venclovas for valuable comments about the manuscript and the software.

