## Supplementary information for "TemStaPro: protein thermostability prediction using sequence representations from protein language models"

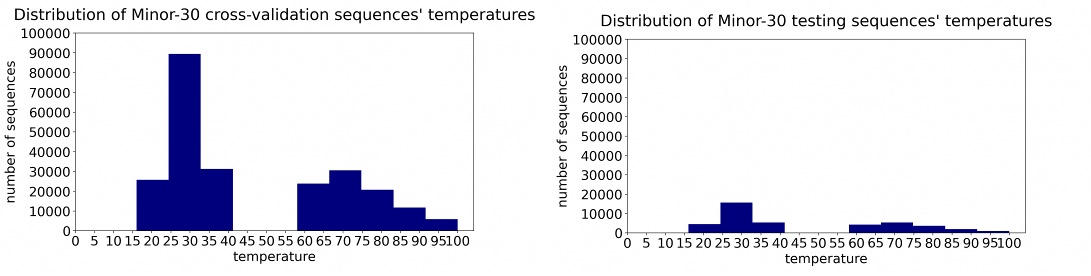

**Figure S1.** Protein sequences distribution regarding to organism's growth temperature in *TemStaPro-Minor-30* cross-validation and testing data subsets.

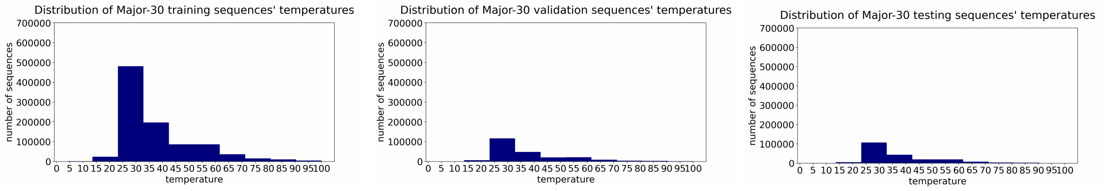

**Figure S2.** Protein sequences distribution regarding to organism's growth temperature in *TemStaPro-Major-30* data subsets.

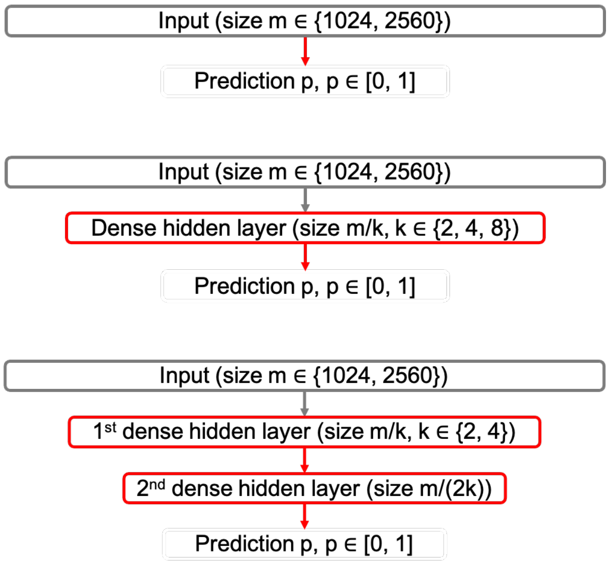

**Figure S3.** Schemes of architectures: single-layer perceptron (upper), a feed-forward neural network model with 1 hidden layer (middle), and a feed-forward neural network model with 2 hidden layers (lower).

**Table S1.** Models that were tested with ESM-2 and ProtT5-XL embeddings as input.

| Model | Number of hidden layers | Size of hidden layers |
|---|---|---|
| C2H2_h1280-640 | 2 | 1280, 640 |
| C2H2_h640-320 | 2 | 640, 320 |
| C2H1_h1280 | 1 | 1280 |
| C2H1_h640 | 1 | 640 |
| C2H1_h320 | 1 | 320 |
| SLP_ESM-2 | 0 | - |

| Model | Number of hidden layers | Size of hidden layers |
|---|---|---|
| C2H2_h512-256 | 2 | 512, 256 |
| C2H2_h256-128 | 2 | 256, 128 |
| C2H1_h512 | 1 | 512 |
| C2H1_h256 | 1 | 256 |
| C2H1_h128 | 1 | 128 |
| SLP_ProtT5-XL | 0 | - |

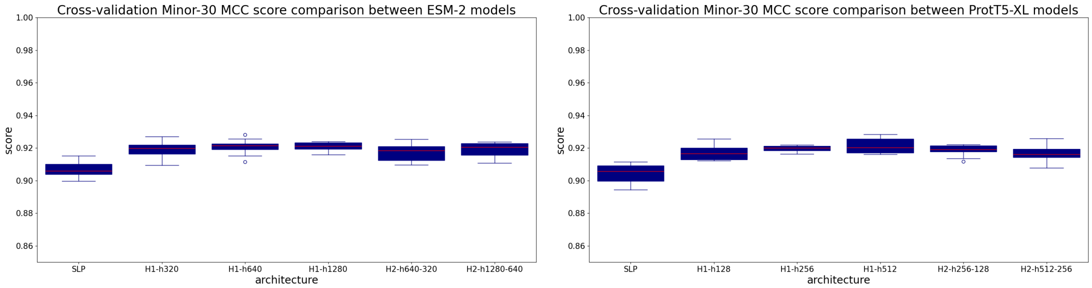

**Figure S4.** Cross-validation MCC scores of different architecture models', which were trained using ESM-2 or ProtT5-XL embeddings of *TemStaPro-Minor-30* set (weight decay equals 0).

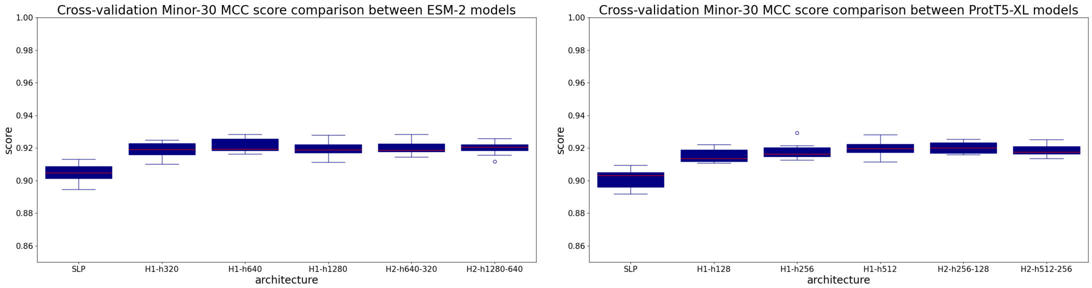

**Figure S5.** Cross-validation MCC scores of different architecture models', which were trained using ESM-2 or ProtT5-XL embeddings of *TemStaPro-Minor-30* set (weight decay $10^{-4}$).

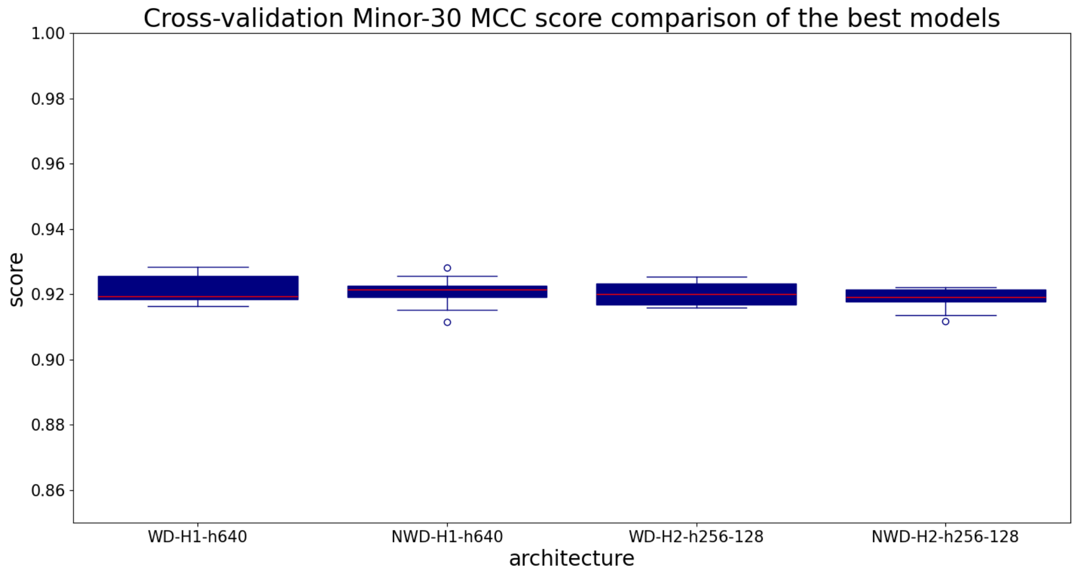

**Figure S6.** Comparison of weight decay effect on the best architectures' (after cross-validation) MCC scores.

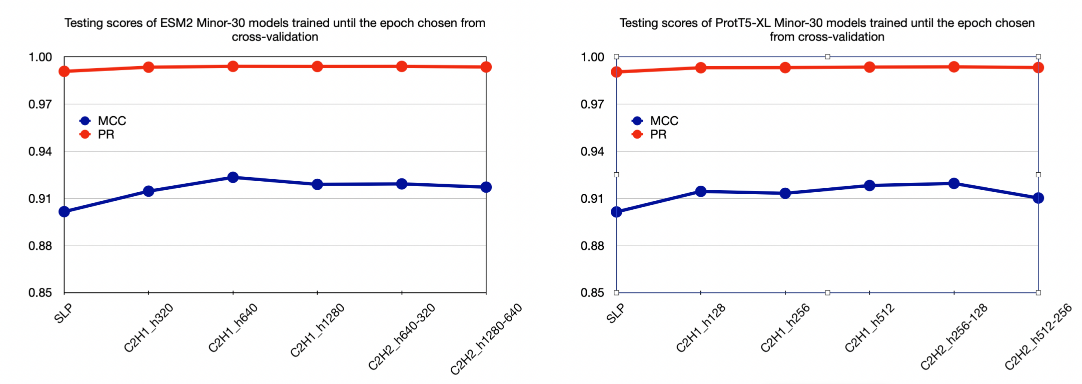

**Figure S7.** Comparison of different architecture models', which were trained using ESM-2 or ProtT5-XL embeddings, MCC and PR AUC scores for *TemStaPro-Minor-30* testing set.

**Table S2.** Model ensembles' scores after testing with the *TemStaPro-Major-30* dataset.

| Model | MCC | Accuracy | Precision | Specificity | Recall | ROC AUC | F1 | PR AUC |
|---|---|---|---|---|---|---|---|---|
| TemStaPro-t80 | 0.515 | 0.947 | 0.295 | 0.947 | 0.956 | 0.990 | 0.450 | 0.768 |
| TemStaPro-t75 | 0.543 | 0.940 | 0.335 | 0.939 | 0.947 | 0.987 | 0.495 | 0.780 |
| TemStaPro-t70 | 0.584 | 0.926 | 0.398 | 0.925 | 0.938 | 0.982 | 0.559 | 0.808 |
| TemStaPro-t65 | 0.610 | 0.919 | 0.449 | 0.919 | 0.922 | 0.978 | 0.604 | 0.820 |
| TemStaPro-t60 | 0.601 | 0.888 | 0.468 | 0.886 | 0.905 | 0.964 | 0.617 | 0.804 |
| TemStaPro-t55 | 0.613 | 0.860 | 0.529 | 0.855 | 0.888 | 0.948 | 0.663 | 0.814 |
| TemStaPro-t50 | 0.638 | 0.856 | 0.605 | 0.854 | 0.865 | 0.939 | 0.712 | 0.838 |
| TemStaPro-t45 | 0.625 | 0.843 | 0.651 | 0.852 | 0.818 | 0.918 | 0.725 | 0.836 |
| TemStaPro-t40 | 0.622 | 0.833 | 0.677 | 0.842 | 0.811 | 0.910 | 0.738 | 0.845 |

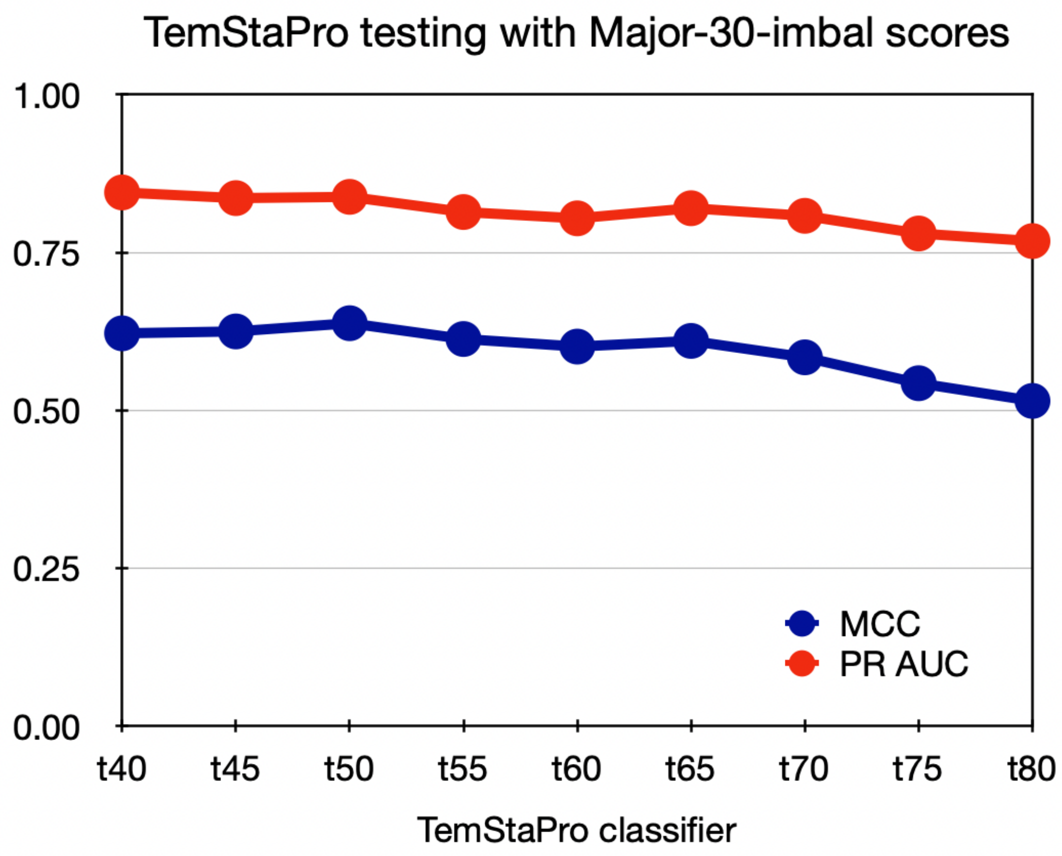

**Figure S8.** Model ensembles' MCC scores after testing with the *TemStaPro-Major-30* dataset.

**Table S3.** Comparison of BertThermo, ProLaTherm, and TemStaPro classifiers' scores using *TemStaPro-Major-30-sample2k* datasets.

| Model | Dataset | MCC | Accuracy | Precision | Specificity | Recall | ROC AUC | F1 | PR AUC |
|---|---|---|---|---|---|---|---|---|---|
| TemStaPro-t80 | bal80 | 0.903 | 0.952 | 0.947 | 0.946 | 0.957 | 0.992 | 0.952 | 0.991 |
| TemStaPro-t75 | bal75 | 0.892 | 0.946 | 0.943 | 0.943 | 0.949 | 0.988 | 0.946 | 0.986 |
| TemStaPro-t70 | bal70 | 0.857 | 0.928 | 0.928 | 0.928 | 0.929 | 0.982 | 0.929 | 0.982 |
| TemStaPro-t65 | bal65 | 0.811 | 0.905 | 0.906 | 0.906 | 0.905 | 0.974 | 0.905 | 0.974 |
| TemStaPro-t60 | bal60 | 0.805 | 0.902 | 0.897 | 0.896 | 0.909 | 0.969 | 0.903 | 0.969 |
| TemStaPro-t55 | bal55 | 0.732 | 0.866 | 0.857 | 0.854 | 0.878 | 0.948 | 0.868 | 0.945 |
| TemStaPro-t50 | bal50 | 0.712 | 0.856 | 0.836 | 0.826 | 0.885 | 0.935 | 0.860 | 0.936 |
| TemStaPro-t45 | bal45 | 0.663 | 0.832 | 0.839 | 0.842 | 0.821 | 0.919 | 0.830 | 0.927 |
| TemStaPro-t40 | bal40 | 0.633 | 0.817 | 0.818 | 0.819 | 0.814 | 0.908 | 0.816 | 0.917 |
| BertThermo | bal80 | 0.700 | 0.848 | 0.810 | 0.787 | 0.908 | 0.847 | 0.856 | 0.882 |
| | bal75 | 0.690 | 0.840 | 0.788 | 0.751 | 0.928 | 0.839 | 0.853 | 0.876 |
| | bal70 | 0.622 | 0.809 | 0.782 | 0.761 | 0.858 | 0.810 | 0.818 | 0.856 |
| | bal65 | 0.624 | 0.811 | 0.792 | 0.778 | 0.845 | 0.811 | 0.818 | 0.857 |
| | bal60 | 0.590 | 0.795 | 0.792 | 0.790 | 0.800 | 0.795 | 0.796 | 0.846 |
| | bal55 | 0.521 | 0.756 | 0.810 | 0.843 | 0.670 | 0.756 | 0.733 | 0.823 |
| | bal50 | 0.446 | 0.718 | 0.775 | 0.821 | 0.615 | 0.718 | 0.686 | 0.791 |
| | bal45 | 0.442 | 0.710 | 0.806 | 0.867 | 0.553 | 0.710 | 0.656 | 0.791 |
| | bal40 | 0.385 | 0.683 | 0.767 | 0.840 | 0.526 | 0.683 | 0.624 | 0.765 |
| ProLaTherm | bal80 | 0.830 | 0.914 | 0.894 | 0.888 | 0.941 | 0.914 | 0.917 | 0.932 |
| | bal75 | 0.797 | 0.898 | 0.891 | 0.889 | 0.908 | 0.899 | 0.899 | 0.923 |
| | bal70 | 0.756 | 0.877 | 0.892 | 0.896 | 0.859 | 0.878 | 0.875 | 0.911 |
| | bal65 | 0.732 | 0.866 | 0.886 | 0.892 | 0.839 | 0.865 | 0.862 | 0.903 |
| | bal60 | 0.704 | 0.847 | 0.919 | 0.933 | 0.761 | 0.847 | 0.833 | 0.900 |
| | bal55 | 0.580 | 0.772 | 0.916 | 0.945 | 0.599 | 0.772 | 0.724 | 0.858 |
| | bal50 | 0.510 | 0.730 | 0.906 | 0.947 | 0.512 | 0.730 | 0.654 | 0.831 |
| | bal45 | 0.482 | 0.705 | 0.932 | 0.968 | 0.442 | 0.705 | 0.600 | 0.827 |
| | bal40 | 0.434 | 0.679 | 0.911 | 0.961 | 0.398 | 0.680 | 0.554 | 0.805 |

| protein_id | position | sequence | length | t40_binary | t40_raw | t45_binary | t45_raw | t50_binary | t50_raw | t55_binary | t55_raw | t60_binary | t60_raw | t65_binary | t65_raw | left_hand_label | right_hand_label | clash |
|---|---|---|---|---|---|---|---|---|---|---|---|---|---|---|---|---|---|---|
| YmeCas12a | - | MSKVNNGF...FVLRNLS | 1362 | 0 | 2.324E-01 | 0 | 2.806E-01 | 0 | 7.563E-02 | 0 | 5.726E-02 | 0 | 1.539E-02 | 0 | 1.216E-02 | <40 | <40 | - |
| SauCas9 | - | MKRNYIL...PQIIKKG | 1053 | 0 | 4.03E-01 | 1 | 5.371E-01 | 0 | 3.865E-01 | 0 | 2.999E-01 | 0 | 8.613E-02 | 0 | 1.718E-02 | <40 | [45-50) | * |
| SpyCas9 | - | MDKKYSI...LSQLGGD | 1368 | 0 | 6.128E-02 | 0 | 4.68E-02 | 0 | 1.984E-02 | 0 | 2.198E-02 | 0 | 9.72E-03 | 0 | 1.842E-03 | <40 | <40 | - |
| CaldoCas9 | - | MRYKIGL...PLQSTRD | 1087 | 1 | 6.278E-01 | 1 | 8.361E-01 | 1 | 7.301E-01 | 1 | 6.187E-01 | 0 | 4.024E-01 | 0 | 2.872E-02 | [55-60) | [55-60) | - |

**Figure S9.** An example tab-separated table that is the output of the (default) global prediction mode of TemStaPro program. The main output of the method is a TSV table with 8 columns: ’protein_id’ - a header taken from the FASTA file of the input protein; ’sequence’ - an amino acid sequence of the protein; ’length’ - a length of the protein’s amino acid sequence; ’t??_binary’ - a binary prediction label for a given temperature threshold (one of the six thresholds is written in the place of question marks) - the label is assigned by rounding the raw prediction (see the next point) at this temperature threshold; ’t??_raw’ - a raw prediction value for a given temperature threshold (real numbers from the interval [0, 1]); ’left_hand_label’ - a label of the highest temperature range, at which the protein was predicted to still be thermostable (possible labels of temperature ranges are: ’<40’, ’[40-45)’, ’[45-50)’, ’[50-55)’, ’[55-60), ’[60-65)’, ’≥65’); ’right_hand_label’ - a label that is interpreted as ’left_hand_label’, yet the label is assigned by reading the outputs starting from the right (possible values of the label coincide with the ’left_hand_label’); ’clash’ - a Boolean identifier, whether a contradiction between the models’ predictions was observed - the expected output is a decreasing sequence of binary predictions if the outputs are read from left to right in the increasing order of the temperature thresholds (expected output is labelled as ’-’ and other cases are assigned ’*’).

| protein_id | position | sequence | length | t40_binary | t40_raw | t45_binary | t45_raw | t50_binary | t50_raw | t55_binary | t55_raw | t60_binary | t60_raw | t65_binary | t65_raw | left_hand_label | right_hand_label | clash |
|---|---|---|---|---|---|---|---|---|---|---|---|---|---|---|---|---|---|---|
| YmeCas12a | - | MSKVNNG...FVLRNLS | 1362 | 0 | 3.663E-01 | 0 | 2.853E-01 | 0 | 1.332E-01 | 0 | 6.786E-02 | 0 | 1.174E-02 | 0 | 1.024E-02 | <40 | <40 | - |
| YmeCas12a | 1 | M | 1 | 0 | 9.439E-02 | 0 | 3.06E-01 | 0 | 7.553E-02 | 0 | 2.346E-03 | 0 | 1.262E-02 | 0 | 4.2E-03 | <40 | <40 | - |
| YmeCas12a | 2 | S | 1 | 0 | 5.779E-05 | 0 | 9.265E-06 | 0 | 1.332E-07 | 0 | 7.257E-07 | 0 | 8.085E-04 | 0 | 8.957E-02 | <40 | <40 | - |
| YmeCas12a | 3 | K | 1 | 1 | 6.523E-01 | 1 | 6.74E-01 | 0 | 2.265E-02 | 0 | 1.152E-01 | 0 | 6.501E-02 | 0 | 9.952E-02 | [45-50) | [45-50) | - |
| YmeCas12a | 4 | V | 1 | 1 | 9.991E-01 | 1 | 9.998E-01 | 0 | 4.998E-01 | 0 | 6.988E-05 | 1 | 7.067E-01 | 0 | 3.767E-01 | [45-50) | [60-65) | * |
| YmeCas12a | 5 | N | 1 | 0 | 2.412E-01 | 0 | 4.262E-01 | 0 | 1.075E-03 | 0 | 5.545E-05 | 0 | 6.789E-08 | 0 | 1.216E-05 | <40 | <40 | - |
| YmeCas12a | 6 | N | 1 | 1 | 9.898E-01 | 1 | 9.989E-01 | 0 | 3.282E-01 | 1 | 5.728E-01 | 0 | 4.837E-01 | 0 | 3.104E-01 | [45-50) | [55-60) | * |
| YmeCas12a | 7 | G | 1 | 1 | 8.477E-01 | 1 | 9.896E-01 | 0 | 6.323E-02 | 0 | 2.215E-02 | 0 | 9.535E-06 | 0 | 4.163E-06 | [45-50) | [45-50) | - |
| YmeCas12a | 1356 | F | 1 | 1 | 9.98E-01 | 1 | 9.991E-01 | 1 | 9.971E-01 | 1 | 9.946E-01 | 1 | 9.872E-01 | 1 | 9.906E-01 | 65<= | 65<= | - |
| YmeCas12a | 1357 | V | 1 | 1 | 9.996E-01 | 1 | 1E+00 | 1 | 9.993E-01 | 1 | 9.982E-01 | 1 | 9.917E-01 | 1 | 9.451E-01 | 65<= | 65<= | - |
| YmeCas12a | 1358 | L | 1 | 0 | 2.587E-03 | 0 | 2.595E-01 | 0 | 4.147E-01 | 0 | 4.249E-01 | 1 | 8.449E-01 | 1 | 9.596E-01 | <40 | 65<= | * |
| YmeCas12a | 1359 | R | 1 | 1 | 7.469E-01 | 1 | 7.959E-01 | 1 | 9.699E-01 | 1 | 7.673E-01 | 1 | 8.938E-01 | 0 | 4.692E-02 | [60-65) | [60-65) | - |
| YmeCas12a | 1360 | N | 1 | 0 | 7.824E-03 | 0 | 3.935E-03 | 0 | 7.495E-03 | 0 | 8.975E-02 | 0 | 8.908E-03 | 0 | 3.43E-01 | <40 | <40 | - |
| YmeCas12a | 1361 | L | 1 | 0 | 3.454E-06 | 0 | 2.247E-06 | 0 | 9.429E-06 | 0 | 1.137E-05 | 0 | 1.454E-03 | 0 | 5.699E-03 | <40 | <40 | - |
| YmeCas12a | 1362 | S | 1 | 0 | 5.388E-04 | 0 | 1.277E-02 | 0 | 8.619E-03 | 0 | 1.917E-01 | 0 | 1.648E-03 | 0 | 5.585E-02 | <40 | <40 | - |

**Figure S10.** An example tab-separated table that is the output of per-residue prediction mode of TemStaPro program.

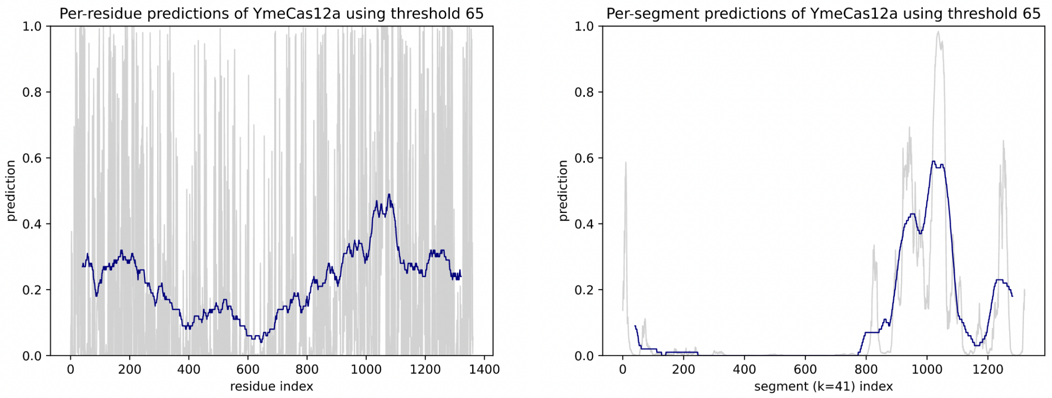

**Figure S11.** An example plot for the output of per-residue mode (left) and per-segment mode with default window size of 41 (right).

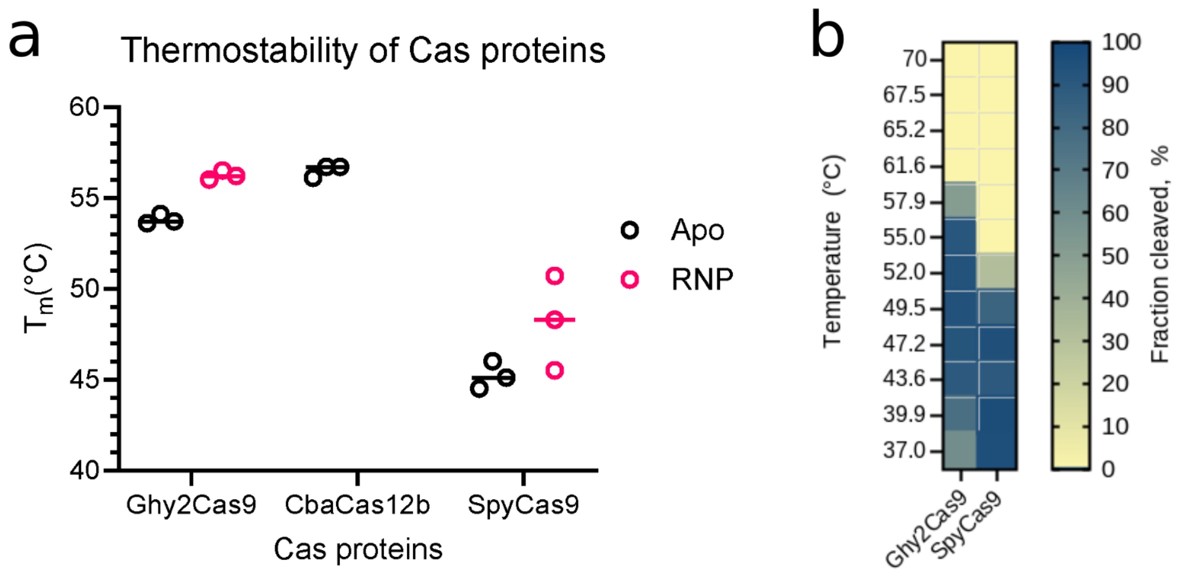

**Figure S12.** **(a)** Thermal stability of Cas proteins without guide RNA (apo) or loaded with sgRNA (Ghy2Cas9 and SpyCas9) (RNP). Protein unfolding was measured using nano differential scanning fluorimetry (nanoDSF) over a temperature range from 20 °C to 80 °C. Fluorescence was monitored as temperature increased at a rate of 1 °C per second. The inflection point of the fluorescent curve is interpreted as the unfolding point of the protein ($T_m$). Data points collected from replicate experiments are plotted as circles, the means are plotted as dashes. **(b)** The double-stranded DNA (dsDNA) cleavage activities of Ghy2Cas9 and SpyCas9 RNPs were measured using in vitro assays containing fluorophore-labeled dsDNA target substrates. Cleaved fragments were quantitated and are represented in a heatmap showing overall activity at temperatures ranging from 37 °C to 70 °C. The intensity of the blue colour indicates the fraction of substrate cleaved.

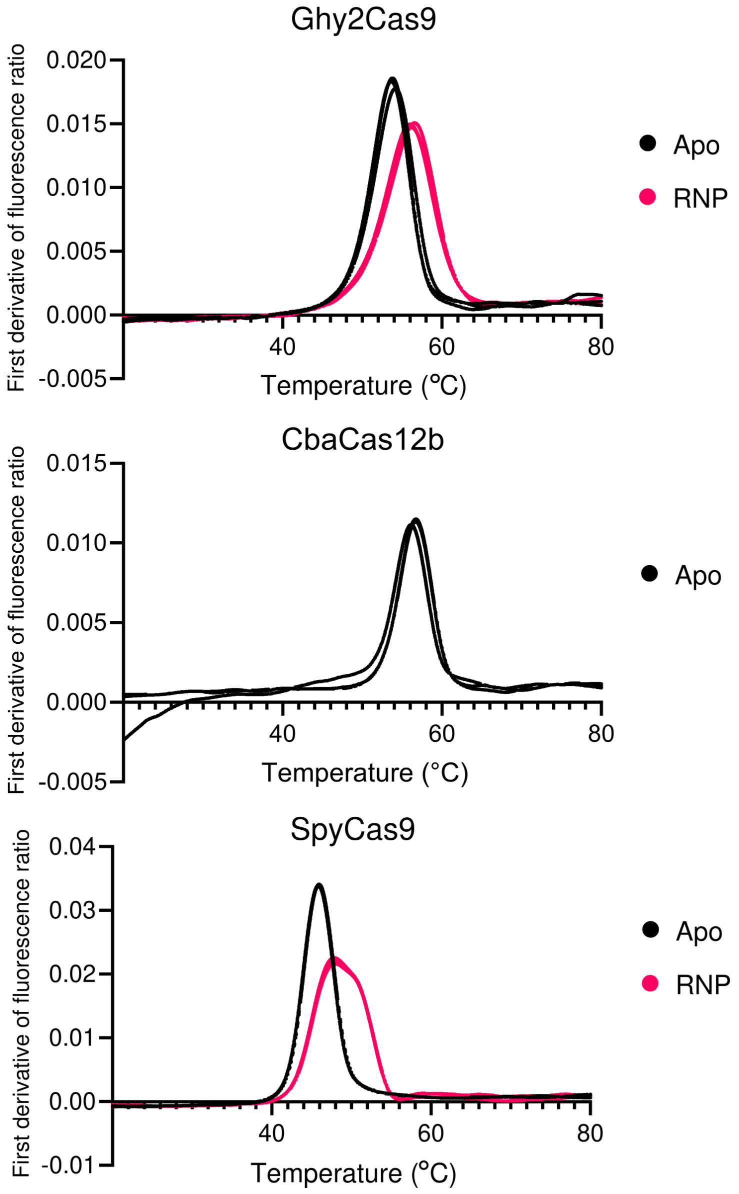

**Figure S13.** Raw traces of nanoDSF assays (*SupplementaryMeltingTemperatureSourceData.xlsx*). The first derivative of the fluorescence ratio was graphed using the integrated fluorescence data at 350 nm and 330 nm by the Prometheus instrument during experiments where the temperature was increased at the rate of 1 °C per second for Cas proteins without guide RNA (apo) or with guide RNA (RNP). The peaks in the first derivative graph are inflection points that correspond to the temperature at which half of the molecules have undergone transition to a new state, which is interpreted as the protein unfolding (denaturing or melting).

**Table S4.** Experimentally characterized and predicted temperatures of C2EP proteins. The full table is in *SupplementaryTableCharacterizedC2EPs.xlsx*.

| Group | Name | Length | Max. active temp | Max. optimal active temperature | Melting point | Reference PubMedID | Prediction left-hand label | Prediction right-hand label | Prediction clash | Difference more than 10 °C[2] |
|---|---|---|---|---|---|---|---|---|---|---|
| Cas12a | YmeCas12a | 1362 | 60 | - | 60 | 27989439 | <40 | <40 | - | Yes |
| Cas12a | CmeCas12a | 1288 | 55 | - | 55 | 27989439 | <40 | <40 | - | Yes |
| Cas12a | RbCas12a | 1247 | 42 | - | 45 | 36012553 | <40 | <40 | - | No |
| Cas12b | AacCas12b | 1129 | 55 | - | - | 27989439 | [45-50) | [45-50) | - | No |
| Cas12b | BrCas12b | 1090 | 63 | - | - | 29127284 | [55-60) | [55-60) | - | No |
| Cas12b | AaCas12b | 1129 | 65 | 59 | - | 30510770 | [50-55) | [50-55) | - | No |
| Cas12b | AacC2c1 | 1129 | 60 | - | - | 27984729 | [55-60) | [55-60) | - | No |
| Cas12b | CbaCas12b | 1120 | - | - | 57 | This study | [60-65) | [60-65) | - | No |
| Cas13a | TccCas13a | 1225 | 70 | - | - | 35763567 | [40-45) | [40-45) | - | Yes |
| Cas13x | mCas13 | 873 | 50 | 37 | - | 34546709 | <40 | <40 | - | No |
| Cas9-C1 | VpaCas9 | 1398 | 48 | - | 43 | 33139742 | <40 | <40 | - | No |
| Cas9-C1 | KhuCas9 | 1309 | 46 | - | 33 | 33139742 | <40 | <40 | - | No |
| Cas9-C1 | FmaCas9 | 1348 | 44 | - | 41 | 33139742 | <40 | [45-50) | * | - |
| Cas9-C1 | EitCas9 | 1330 | 44 | - | 37 | 33139742 | <40 | <40 | - | No |
| Cas9-C1 | Sag1Cas9 | 1384 | 44 | - | 40 | 33139742 | <40 | <40 | - | No |
| Cas9-C1 | Sag2Cas9 | 1377 | 41 | - | 41 | 33139742 | <40 | <40 | - | No |
| Cas9-C1 | SdyCas9 | 1371 | 48 | - | 45 | 33139742 | <40 | <40 | - | No |
| Cas9-C1 | SmuCas9 | 1345 | 48 | - | 45 | 33139742 | <40 | <40 | - | No |
| Cas9-C1 | SpyCas9 | 1368 | 50 | 42 | 47 | 28146359 | <40 | <40 | - | No |
| Cas9-C2 | Cme3Cas9 | 1124 | 54 | - | 50 | 33139742 | [50-55) | [50-55) | - | No |
| Cas9-C2 | Ghh1Cas9 | 1094 | 37 | - | 41 | 33139742 | <40 | <40 | - | No |
| Cas9-C2 | CgaCas9 | 1403 | 37 | - | 36 | 33139742 | <40 | <40 | - | No |
| Cas9-C2 | Cca1Cas9 | 1430 | 44 | - | 41 | 33139742 | <40 | <40 | - | No |
| Cas9-C2 | Cme1Cas9 | 1399 | 41 | - | 47 | 33139742 | <40 | <40 | - | No |
| Cas9-C2 | OrhCas9 | 1535 | 44 | - | 43 | 33139742 | <40 | <40 | - | No |
| Cas9-C2 | WviCas9 | 1440 | 48 | - | 43 | 33139742 | <40 | <40 | - | No |
| Cas9-C2 | Ghy2Cas9 | 1358 | 58 | 55 | 57 | This study | [55-60) | [55-60) | - | No |
| Cas9-C2 | IgnaviCas9 | 1244 | 90 | - | - | 31659048 | [55-60) | [55-60) | - | Yes |
| Cas9-C3 | TmoCas9 | 1049 | 46 | 37 | 48 | 33139742 | <40 | <40 | - | No |
| Cas9-C3 | Ghy3Cas9[1] | 972 | 46 | - | - | 33139742 | [55-60) | [55-60) | - | No |
| Cas9-C3 | CaldoCas9 | 1087 | 60 | - | - | 33953310 | [55-60) | [55-60) | - | No |
| Cas9-C3 | GeoCas9 | 1087 | 65 | - | - | 29127284 | [55-60) | [55-60) | - | No |
| Cas9-C3 | ThermoCas9 | 1082 | 60 | - | - | 29162801 | [55-60) | [55-60) | - | No |
| Cas9-C4 | Sth1ACas9 | 1122 | 57 | - | 50 | 33139742 | <40 | <40 | - | No |
| Cas9-C4 | SauCas9 | 1053 | 50 | - | 50 | 33139742 | <40 | [45-50) | * | - |
| Cas9-C4 | MgaCas9 | 1269 | 46 | 44 | 47 | 33139742 | <40 | <40 | - | No |
| Cas9-C4 | SsaCas9 | 1127 | 50 | - | 41 | 33139742 | <40 | <40 | - | No |
| Cas9-C4 | SsiCas9 | 1122 | 48 | - | 44 | 33139742 | <40 | <40 | - | No |
| Cas9-C4 | SsuCas9 | 1122 | 44 | - | 42 | 33139742 | <40 | <40 | - | No |
| Cas9-C5 | AceCas9 | 1138 | 60 | 50 | - | 28277645 | <40 | <40 | - | No |
| Cas9-C7 | NsaCas9 | 1137 | 64 | - | 67 | 33139742 | [60-65) | [60-65) | - | No |

[1] - mislabeling in Fig. 4A of 33139742 instead of Ghy2, should be Ghy3;
[2] - prediction differs more than 10 °C from max active/optimal temperature and protein melting point.
